# “I’d like to think I’d be able to spot one if I saw one”: How science journalists navigate predatory journals

**DOI:** 10.1101/2024.07.24.604934

**Authors:** Alice Fleerackers, Laura L. Moorhead, Juan Pablo Alperin

## Abstract

Predatory journals—or journals that prioritize profits over editorial and publication best practices—are becoming more common, raising concerns about the integrity of the scholarly record. Such journals also pose a threat for the integrity of science journalism, as journalists may unwillingly report on low quality or even highly flawed studies published in these venues. This study sheds light on how journalists navigate this challenging publishing landscape through a qualitative analysis of interviews with 23 health, science, and environmental journalists about their perceptions of predatory journals and strategies for ensuring the journals they report on are trustworthy. We find that journalists have relatively limited awareness and/or concern about predatory journals. Much of this attitude is due to confidence in their established practices for avoiding problematic research, which largely centre on perceptions of journal prestige, reputation, and familiarity, as well as writing quality and professionalism. Most express limited awareness of how their trust heuristics may discourage them from reporting on smaller, newer, and open access journals, especially those based in the Global South. We discuss implications for the accuracy and diversity of the science news that reaches the public.

---

In 2023, more than 10,000 journal articles were retracted, setting an all-time record (Van Noorden, 2023). While the sheer volume of articles retracted in 2023 is alarming, it is not an isolated event, but rather a continuation of a longer-term trend with far-reaching effects (*Ibid*.). Notably, it can take years for problematic research to be retracted (if it is retracted at all), and many retracted papers continue to receive citations and news coverage (Bolland et al., 2022).

Fueled by an academic system that rewards researchers based on their publication records, some researchers fake data, report analyses they never conducted, and publish overblown or even doctored results (H. Shen, 2020; West & Bergstrom, 2021). *Paper mills* churn out fraudulent articles on demand for exorbitant fees (Else & Van Noorden, 2021). Meanwhile, *predatory journals* “prioritize self-interest at the expense of scholarship,” publishing articles for profit, with little or no adherence to best editorial and publication practices (Grudniewicz et al., 2019, p. 211). While such journals may claim to have a robust peer review process, many provide little or no such review in practice (IAP, 2022).

All this has the potential to compromise the accuracy of the science and health news that ultimately reaches the public, as journalists consistently rely on journal articles as a primary source for information in their reporting (Wihbey, 2017). They rarely report or follow up when an article is retracted, and, as a result, unsupported findings can gain wide media coverage before they are eventually debunked—if they are debunked at all (Rada, 2007; Santos-d’Amorim et al., 2021). When later studies inevitably challenge the scientific “breakthroughs” presented in the news, frustration can ensue, negatively impacting citizens’ willingness to make evidence-based health decisions, their receptivity to future health recommendations, and their trust in journalism (Nagler, 2014; Nagler et al., 2022, 2023).

Predatory publishing is not a black-and-white phenomenon but rather one with many shades of gray, as even some “legitimate” journals use predatory practices at times (Amsen, 2024; IAP, 2022; Nicholas et al., 2023), further complicating journalists’ task of distinguishing between problematic and trustworthy science. Indeed, some have argued that the high Article Processing Costs (APCs) or aggressive solicitation practices used by some “legitimate” journals and publishers could arguably be considered predatory (Linacre, 2022). Others note that even predatory journals may publish high-quality research and argue against assessing research based on the “container” it comes in rather than its “contents” (Eve & Priego, 2017) Moreover, commonly used strategies for identifying predatory journals have been critiqued for being unscientific and biased against OA journals and those based in the Global South (Amsen, 2024; Krawczyk & Kulczycki, 2021).

These nuances and complexities can make it difficult to determine whether a journal is trustworthy (IAP, 2022). It is likely that doing so is especially challenging for journalists without advanced science training. However, scholarship examining journalists’ awareness or use of predatory journals is extremely limited, as are studies of their practices for navigating this expanding publishing landscape. This study contributes to filling this gap through an analysis of interviews with 23 health, science, and environmental journalists about their perceptions and use of predatory journals.

## Literature review

### Journalists’ perceptions and use of predatory journals

A small survey of US health and science journalists provides preliminary evidence that some journalists worry about encountering predatory journals and have developed strategies, albeit largely superficial, for ensuring they avoid using them (Schultz, 2023). Of the 82 journalists in the study, about three quarters were familiar with predatory journals and almost all of those who were familiar were highly or somewhat concerned about them. Journalists’ understanding of predatory journals and publishers primarily centred on the financial motivations of these outlets, although a handful of participants noted other factors, such as their lack of quality control. To avoid using research from predatory journals, journalists reported examining the journal’s website, seeking input from researchers, or referencing online lists of predatory publishers. They considered factors such as whether the journal website included grammar or spelling errors, was transparent about the journal’s publishing practices, and included articles written by other, known researchers.

Beyond these early findings, existing evidence is extremely limited. However, prior scholarship on journalists’ use of research more broadly has relevance to their engagement with predatory journals. Specifically, studies have found that journalists preferentially report on elite, high impact journals such as *Science* and *Nature* (Hansen, 1994; MacLaughlin et al., 2018; Moorhead et al., 2021), as well as on articles that have been promoted by a press release (Bray, 2019; McKinnon et al., 2019). They rely heavily on interviews with researchers to decide which studies to report on, vet their quality, and frame their results and implications (Albæk, 2011; Fleerackers et al., 2022; Gesualdo et al., 2020). Journalists also tend to defer to the peer review process rather than verifying research themselves, placing their trust in journals and the scientists who review for them (Fleerackers & Nguyen, 2024). This reliance on scientists has often been critiqued, as it may encourage “science cheerleading” and impede journalists’ ability to report critically on new studies (Blum, 2021; Figdor, 2017). Indeed, media coverage of research often presents scientific findings as established facts, with limited discussion of associated caveats, biases, or problematic implications that may compromise the public interest (Dumas-Mallet et al., 2018; Guenther et al., 2019; Matthias et al., 2020). While scholarship is lacking, it is possible that journalists’ relatively uncritical trust in science may lead them to report on predatory journals, some of which closely resemble well-established journals and, at times, rank higher than them in online search results (IAP, 2022; Siler et al., 2021). Thus, there is a need to explore the professional practices that science journalists rely on to navigate predatory publishing.

### Predatory journals and inequities in scholarly communication and science journalism

Journalists’ relationship to predatory journals not only has implications for the accuracy of science news, but also its representativeness. Specifically, a wariness of predatory publishing among journalists may encourage them to avoid research published in smaller, lesser known journals, even though such journals may have a more robust peer review process in place than their elite competitors (Oransky, 2022) and likely publish research that is more relevant to audiences’ geographic and cultural contexts (Fleerackers & Nguyen, 2024). Additionally, avoiding research published in lesser-known journals can exclude early career scholars and those working outside the realm of “highly visible scientist,” “celebrity scientist,” or media pundit (Fahy & Lewenstein, 2021). Moreover, Global South scholars are often more vulnerable to publishing in predatory journals due to an “increasingly globalized and homogenized view of academic excellence based on ‘where’ and how often one publishes,” which puts pressures on these scholars to demonstrate their international impact (C. Shen & Björk, 2015, p. 14). As a result, the perceived threat of predatory publishing may disproportionately raise suspicion about journals and researchers from the Global South, whose articles already receive greater scrutiny within academia and journalism (Abdill et al., 2020; Nguyen & Tran, 2019). Journalists’ tendency to equate grammar and spelling errors with predatory publishing (Schultz, 2023) may further enhance this potential bias, discouraging journalists from reporting on research published by authors whose first language is not English.

More broadly, predatory journals’ pay-to-publish model means these journals are often conflated with open access (OA) publishing (Krawczyk & Kulczycki, 2021), which may discourage journalists from reporting on OA journals. Indeed, while research on journalists’ perception of OA is limited (Fleerackers et al., 2024), preliminary studies suggest that at least some journalists are “more suspicious of OA journals, believing they lacked a credible review process” (Van Witsen & Takahashi, 2021, p. 10, also Schultz, 2023). As many OA journals are based in the Global South, these suspicions may further entrench existing imbalances in science news coverage (Fleerackers & Nguyen, 2024; Nguyen & Tran, 2019).

In other words, predatory journals are not just an integrity problem; they “are a geopolitical problem because the geopolitical peripheries of science are much more often harmed by them than the center” (Krawczyk & Kulczycki, 2021, p. 1; also C. Shen & Björk, 2015). Yet the ways in which journalists’ perceptions of predatory journals may be impacted by, and, in turn, impact this geopolitical problem are not yet well understood. This research begins to shed light on this question by addressing the following research questions:

RQ1. How do journalists perceive predatory journals?

RQ2. How do journalists decide whether a journal is trustworthy?

RQ3: What are the potential implications for these perceptions and practices in terms of the scholarship that is featured in the news?

## Methods

Informed by a pragmatist paradigm, we conducted a qualitative interview study using interpretive description. This approach was selected because of its usefulness for developing insights with practical relevance (Creswell & Poth, 2017; Thorne et al., 1997). The study is part of a larger project examining how journalists understand and respond to journals’ adherence to scholarly integrity standards. The protocol was approved by the Ethics Review Boards of San Francisco State University (#2023–157) and Simon Fraser University (#30001768).

We recruited journalists to participate in semi-structured interviews by advertising calls for participants through professional organizations and social media groups for science journalists and communicators (e.g., the Public Communication of Science and Technology Network’s listserv, Binders Full of Science Writers Facebook Group). Interviews took place via Zoom and were recorded, transcribed, and de-identified for analysis. We recruited and interviewed journalists until our analysis suggested we had reached *information sufficiency,* i.e., we had enough data to comprehensively answer our research questions (LaDonna et al., 2021; Vasileiou et al., 2018).

The final sample comprised 23 health, science, and environmental journalists from six countries (Canada, Denmark, England, Mexico, US, and Switzerland). Most participants were US-based (*N*=14) freelancers (*N*=14) and had 10 or more years of journalism experience (*N*=14). All journalists had a university degree, most typically in science writing/communication (*N=*8), journalism (*N*=6), and/or a STEM field (*N*=9). Four journalists did not have a postgraduate degree (e.g., master’s or PhD). Interview questions centred on journalists’ professional backgrounds, motivations for using research, strategies for finding and vetting studies and journals, and their perceptions of predatory journals and other scholarly communications topics (e.g., retractions, preprints, research integrity). Journalists also completed a walkthrough of two sample papers published in two unnamed journals. The journalists described the steps they would typically take to verify that the research could be trusted. A complete interview protocol is available online (Moorhead & Fleerackers, 2024).

We analyzed the interview transcripts collaboratively through reflexive thematic analysis, using our own subjectivity and “dual processes of immersion or depth of engagement, and distancing” to develop themes relevant to our research questions (Braun & Clarke, 2022, p. 9).

AF and LLM began by familiarizing themselves with the interviews and taking independent notes about potential codes and themes via Google Docs, guided by a broad research objective of understanding journalists’ perceptions and practices for assessing journal and research trustworthiness. They then met to discuss areas of synergy and divergence before conducting further independent coding. This second round of coding prompted AF to recognize a recurring pattern within the sections of the interviews where journalists discussed predatory journals, which guided her to draft a preliminary set of themes and research questions focused on the topic. This draft was then shared with all authors, who scrutinized, questioned, and added to the emergent themes during follow up meetings and via asynchronous editing. Guided by this input, AF performed the final coding using NVivo 14. She focused this analysis on sections of the transcripts that related specifically to predatory journals and journal trustworthiness but also considered additional sections to contextualize themes.

In line with reflexive thematic analysis, the research team’s expertise and interest in journalism, open science, journal integrity, and equity and diversity provided a distinct lens that impacted the analysis and interpretation of the data. Specifically, all three authors believe in the value of OA publishing and are motivated to ensure a diversity of research outputs are recognized within both scholarly communication and journalism. These beliefs prompted us to interrogate potential implications for both OA and epistemic diversity when analyzing how journalists assessed research trustworthiness.

## Results

### Predatory journals: A problem in theory but not in practice

When asked whether they worried about predatory journals, journalists noted that they were problematic in theory but didn’t see them as an obstacle—or, in some cases, even a concern—in their own work. Although one journalist believed that “any journalist has probably encountered them or pitched stories by them” [J1], most shared a conviction that they never “really come across any predatory journals” and so “it’s not really a problem, or hasn’t been a problem, for me” [J3]. To journalists, predatory journals were something they worried about “in a philosophical sense, but I feel like I don’t run across them that often, honestly…I’d like to think I’d be able to spot one if I saw one” [J4].

A few journalists appeared to have limited or no knowledge of predatory publishing, admitting that they “haven’t really heard of that” [J5] or that they were more interested in vetting study quality and hadn’t “really put a lot of thought into the journals themselves” [J9]. Some required an explanation of the term *predatory journal* before they could answer our interview questions. That is, for these journalists, their unworried attitude was due to a lack of awareness, not a lack of concern about problematic research.

One journalist, with a PhD in a STEM field and a previous career in academia, voiced a concern about the difficulty journalists face in deciding if an unfamiliar journal is predatory: “I don’t really know how you can judge as a complete outsider for a journal you don’t know…It’s a problem” [J18]. The journalist added that journal status is not stationary: “Some publications I used to consider credible five or six years ago, when I was doing my research, are now…moving in a real predatory direction with the quality criteria decreasing and peer review decreasing” [J18]. Similarly, J4 noted that it can be “hard to tell what the line is between predatory and not predatory. Arguably, a lot of practices of a lot of the big journals that we read, I think, are predatory.” These nuanced perspectives were relatively rare, however, and may be linked to the professional backgrounds of these journalists, both of whom had advanced STEM training.

More commonly, participants expressed confidence that they would never cover research from a predatory journal, because they had honed their ability to discern credible from problematic research through years of experience. As J5 explained, “at this point in my career, having done this for, what, over eight years, I’ve just developed a set of standards internally, that I, I trust my own processes, and I trust my gut.” Other journalists similarly pointed to the highly internalized nature of this “set of standards,” which appeared to operate like intuition. They made comments such as, “I mean you can just tell when, like, a manuscript has just been slapped together, right?” [J21], and “everyone knows what is a wrong [journal] and what is a more ethical [journal]” [J20]. As J6 explained:

> I don’t really worry about [encountering a predatory journal]. I think when you, when you get used to looking for good research, and, you know, quality research with an interesting finding, it’s just not going—you’re not going to find that in a predatory journal.

### “Reputable” and “familiar” as proxies for “trustworthy”

When it came to discerning predatory from credible journals, the journalists we interviewed voiced many of the same biases that have been documented in previous literature. Specifically, these journalists often placed their trust in high-impact, prestigious journals such as *Nature* or *Science,* as well as known publishers such as Sage or Taylor & Francis. Journalists noted that they would never encounter predatory journals because, “generally, the ones that I consult are pretty well established” [J13]. Similarly, predatory journals were not a problem for journalists such as J12, who explained: “I don’t need to go that far to get information…I don’t look beyond the—you know, the really, the mainstream journals.”

As discussed above, this ability to distinguish “mainstream” and “well established” journals from those that are less well regarded was often referred to in a taken for granted way, as if it were a skill every science journalist possesses and practices intuitively. However, journalists also described using scholarly rankings and metrics as indicators of whether a journal was reputable, and, by extension, trustworthy. For example, J4 reported that they “evaluate ‘What is the impact factor?’” when vetting an unknown journal, while J22 sometimes considered “how the journal ranks compared to other journals.”

Perceptions of prestige or reputation were often closely intertwined with journal familiarity. Specifically, journalists repeatedly described applying greater scrutiny when encountering journals they’d never heard of, while those they used frequently (or saw their colleagues report on) were seen as more trustworthy. J7 said that “if I don’t recognize a journal name, I will definitely go to their page and see what type of stuff they’re publishing,” while J9 reflected that “I guess I maybe have seen a journal here and there that I’ve never seen before, and maybe that’s when I more skeptical.” Journal (un)familiarity was often the first “red flag” journalists pointed to when asked how they would determine whether a journal was predatory.

Journalists varied in the degree to which they relied on reputation and familiarity as markers of trustworthiness. For example, while examining one of the sample papers during the walkthrough portion of the interview, J7 noted, “So, I see this [study] was in *Science*, so, kind of automatically, I’m like, ‘Cool, this is legit,’ because *Science* wouldn’t publish anything bad.” At the extreme, journalists claimed that they “wouldn’t ever report on something from a journal that I had never heard of,” explaining:

> I vet research largely by the publication that it’s in. So, if it’s in a weird publication, I sort of don’t know whether to trust it or not. I usually just don’t, because I don’t, I sort of don’t know. I’m like, well, it would be in a better publication if it were trustworthy [J14].

However, some journalists were more cautious, noting that even high-impact, reputable journals such as *The Lancet* sometimes let problematic research “slip through” [J1] or publish “all kinds of crap” [J22]. Still, even these journalists described the reputation of the journal or the publisher as “shorthand” or a “very hard and fast rule” [J4] that could be used to get at least a preliminary sense of whether a particular study was rigorous. The reputations of the journal and publisher were referred to in an offhand way, as if they were “obvious signifiers” [J6] of research quality whose validity and usefulness required little or no explanation.

### Publication and editorial practices

Familiarity and prestige were not the only markers journalists used to assess a journal’s trustworthiness. Both when describing the qualities of predatory journals and of those they deemed credible, journalists also mentioned criteria related to publication and editorial practices.

Chief among these was the quality or presence of the peer review process in place at the journal. Journalists often defined predatory journals on this basis, describing them as “journals that basically, like, they have no filtering process” [J21], or stating that, “Well, they don’t do this peer review or they say that they do peer review, but at the end, the data is not confident” [J20]. For some journalists, these perspectives were linked to an unwavering trust in the peer review process—especially at top-tier journals. As J3 expressed, “anything that’s been published like that in a peer reviewed journal is going to be obviously true and factual and correct.” Similarly, J12 exclusively covered “mainstream” journals because “the main journals, they do have a good reputation for [peer review]. Their reputation’s on the line when they—if they don’t, if they publish something that’s not good.”

Closely related to having a rigorous peer review process, journalists found it important that journals were selective in what they published. Stringent acceptance rates suggested a journal was trustworthy, because “that tells you, ‘Is it a predatory journal or does it pick and choose?” [J21]. Similarly, upon being presented an example of a journal acceptance rate during the walkthrough portion of their interview, J19 commented that, “It looks like it is a selective journal, which is nice. It doesn’t appear to be a predatory journal.”

In addition, journalists expected credible journals to put research articles through a thorough editing or proofreading process. As a result, journals containing articles that were not clearly written, or contained grammatical or spelling errors, immediately sparked suspicion. For example, J19 stated that “most scientific journal articles that I read are in the English language. If no one has even bothered to edit the English to make it comprehensible, then that’s a red flag.” Similarly, during their walkthrough, J21 became skeptical after noting some typos in an article:

> And then some of the words under “Abstract,” like, just the sentences, it’s a bit awkward. Like, the way it’s written, it’s a bit awkward. So right away to me that says, you know, they didn’t have a process in place where someone actually edited these things. So right off the bat to me that’s a red flag.

### Funding models: The complexity of open access publishing

Finally, some journalists described a journal’s funding model as an important clue for ascertaining whether it was credible. They explained that predatory journals operated on “a sort of profit-driven motive” [J22], charging authors “a high cost of pay to play” [J23] to publish in them. Similarly, J11 said that, “obviously, there are excellent journals of all kinds, but knowing what their source of funding is or their business model is could be helpful” for determining their trustworthiness. Interestingly, some journalists who were weary of such for-profit publishing models also acknowledged that “publishing is a business, and so that’s just the reality of things. Like typically, the most elite journals, you have to pay” [J5].

This focus on funding models raised tensions when it came to OA journals. Many journalists saw OA as the ethical and fair approach for sharing publicly financed science and for expanding perspectives and voices in research. On a practical level, these journals were seen in a positive light by journalists who faced challenges in accessing (non-OA) research. For example, J19 had “recently lost” journal access through their former institution and had turned to OA journals as a “lifeline” for circumventing paywalls. They described “trying to make the open access research” work, but still experienced the situation as “a real barrier” as they were only able to “access only about a third of the information” they needed [J19]. In some cases, OA status encouraged journalists to look favorably on the journal. For example, when J18 noticed that the journal they were examining during their walkthrough was OA, they stated, “So, it’s open access. I’m hooked.”

Yet other journalists considered OA a space fraught with editorial risk and, increasingly, a front for publishers with predatory and exploitative motives. For instance, J5 explained that they never report on OA journals but admitted that “I feel self-conscious about saying that because I’m sure that there are open access publishers that are great.” Interestingly, this journalist became increasingly aware of the impacts of their biases throughout the interview, noting towards the end that they “feel like I prejudged the open access thing.”

Another journalist, a proponent of universal OA for research, expressed frustration about how some legitimate OA journals had started adopting predatory publishing practices. The journalist explained:

> Open access has to be the norm…but the money has to come from somewhere. The problem with [name of a gold open access publisher]—it started as a fantastic initiative; it’s a very noble goal and there are many [of their] journals that I value as high quality— but there are many others in that group that I would start classifying as predatory. There is a big gap, a space for predatory journals—and they know it [J20].

For this journalist, the risks of OA and predatory journals were intertwined with the cost of academic publishing, particularly APCs. For instance, the journalist worried that APCs kept researchers from publishing in top-tier journals, encouraging them to look for cheaper or no-cost alternatives and potentially publishing in lower quality or even fraudulent journals. The journalist suggested that a fear of these publications could encourage journalists to avoid OA journals altogether, further limiting public access to knowledge. The journalist also believed that APCs had created an access divide of “haves and have-nots,” especially for those in developing countries or early in their careers [J19].

Other journalists were ambivalent about OA and did not consider this when deciding whether to trust a journal. For instance, J22 labeled themselves “neutral” in the access debate, calling OA “a different way to get money from the process.” As a former news editor affiliated with a scientific journal, they found OA to be “such a complicated question” and one that “is rarely treated with the nuance it deserves.”

### The threat of the predatory and its impact on source diversity

Although journalists considered multiple factors when discerning whether a journal was trustworthy, journal prestige and familiarity were by far the most relied on “safety measure[s]” [J12]. While this reliance on journal reputation and familiarity has been documented before, our interviews suggest that this tendency may be becoming even more entrenched as journalists navigate growing concerns about predatory publishing and related research integrity issues.

As J1 explained, “with the rise of a lot of like, predatory journals, and like, you know, just pay and get an article published. Like, I find that, in general, you know, we’re a lot choosier.” Similarly, journalists voiced concerns about issues such as academic fraud, hyped or untransparent research reporting, and the rise of generative AI in academic publishing. These concerns, in turn, encouraged more conservative research selection practices. For instance, J8 explained:

> …with all of these AI and technologies emerging, it’s kind of hard to tell if the information is trustworthy or not. So, because of that, I only rely on trustworthy sources. I don’t go by information in all of the places, but I only report to those that are trustworthy or recommended by my other journalist fellows or, as I see, many of the other people have used those [sources] in publication.

Similarly, J1 recalled encountering many journals “that were publishing clearly things that were not at all evidence-based” while reporting on COVID-19 vaccine science during the pandemic. This awareness that low-quality or problematic research could be published in peer reviewed journals permanently altered their approach to vetting research: “I became much, much more skeptical of journals that…didn’t have a track record that I kind of knew, like they had this new name that you never really heard of” [J1].

Importantly, for some journalists, concerns about research integrity had the opposite effect, discouraging them from relying on journal reputation as an indicator of quality. For instance, J22 remarked that, “yes, I worry about predatory journals, but, honestly, I also worry about hype from completely legitimate journals.” These concerns motivated this journalist to take a highly critical stance when vetting research—regardless of the journal it was published in. Similarly, J16 recounted learning about a hyped and, ultimately, highly flawed research article earlier in their career, which led them to conclude that “even if it’s the most prominent researcher in the most prominent journal, I’m gonna be skeptical.” However, this critical stance was far from universal.

Although journalists repeatedly acknowledged their reliance on journal reputation and familiarity as ways to avoid problematic and predatory science, many were unconcerned or unaware of how doing so might impact the diversity of the research they covered. J7 reflected that, “I think that there are small journals that are unreported that have very valid research in them.” However, this reflection appeared to be prompted by a question asked earlier in the interview, as the journalist added that, “maybe this goes back to your question about how I deem something as being, like, reliable that I didn’t think about” [J7].

Similarly, few journalists acknowledged that their tendency to equate “shoddy” or unprofessional writing to low-quality scholarship may bias them towards covering research from English-speaking countries or scholars or saw this potential bias as problematic. As one journalist remarked when examining a research article during their walkthrough,

> Yeah, I mean, right off the bat I noticed a few typos, which is always a red flag for me…[but] it seems like everybody [who authored this paper] is local to the US, so I’m not sure why there’s these typos here [J21].

A few journalists, however, did express concern about the biases embedded in their strategies for avoiding problematic research. For instance, J17 reflected on the usefulness, but also the elitism, of using a journal’s article acceptance rate to weed out predatory journals:

> This article acceptance rate is—it’s funny because in one way it smacks of elitism— “we’re a better journal because we accept fewer articles”—but it is kind of a gauge. Literally, in my email inbox right now, I’ve got this correspondence with a scientist who—he used ChatGPT to basically make up an article and submit it to a predatory journal that was soliciting information from him. They accepted it without question and essentially were going to publish it until we pointed it out…I’m pretty sure that that journal, for instance, accepts 100 percent of its articles. It doesn’t even reject an AI-written article. So, although this smacks of elitism, it actually is a useful thing.

Similarly, one journalist remarked that vetting publications based on writing quality could be problematic from a source diversity perspective, although they were at a loss of how to address this bias:

> I do a lot of work in agriculture and a lot of agricultural research is done in developing countries. I am not always familiar with [the] journals, but I’m afraid that some could be predatory and I don’t always know it. And sometimes I can tell by the quality of the work, but again, [in] developing countries, the quality of the work may not appear to be as great because people have multiple barriers to publishing [J19].

This relative lack of concern about research diversity persisted for some journalists even after we adapted our interview protocol to include a question about this potential bias. For example, when asked whether they worried that their practices might “limit the research or voices and perspectives included in research and ultimately in reporting,” J21 responded, “No, it’s not a concern.” Others were more ambivalent or pivoted, taking the conversation in a different direction. Still others expressed explicit concern but were at a loss for how to address the problem: “I really gotta pick, like, the biggest, most impactful [study to cover], which often ends up being in the bigger journals…I think part of it’s the nature of my job” [J23].

## Discussion

This study sought to understand how health, science, and environmental journalists perceive predatory journals, how they ensure the journals they report on are credible and trustworthy, and the implications of these perceptions and practices for the diversity of the research that makes the news. Collectively, the findings suggest that these specialized journalists largely believe predatory journals are a problem for their peers, or a problem in theory, but not one they would ever fall for themselves. Many have relatively limited awareness and understanding of these journals and feel confident that they can avoid research published in them by applying internalized strategies for evaluating a journal’s “trustworthiness.” These strategies rely heavily on assessments of a journal’s reputation and familiarity—as well as the perceived professionalism of the articles published within it—and are often applied in an intuitive, taken-for-granted way.

Journalists’ relatively unconcerned attitudes about predatory journals comes with risks, as identifying these journals is notoriously difficult. While some journals are obviously of low quality, others “hijack” established and reputable journals, creating “illegitimate clones” that can appear earlier in search results than the original journal (Siler et al., 2021, p. 564). Similarly, some predatory journals use names that closely resemble those of known journals or are indexed by well-known services such as PubMed or Scopus (IAP, 2022), which could mislead deadline-driven journalists looking for story ideas or evidence and sourcing for an article they are writing. Even academics can be misled by predatory tactics, with as many as 14% of researchers in one global survey reporting they had participated in a predatory conference or published in a predatory journal and another 10% stating that they were unsure whether they had done so (IAP, 2022). It is possible that journalists, too, unknowingly cover research in problematic journals, compromising the accuracy of the health and science information that reaches the public.

Moreover, there is a growing consensus that most journals do not fall neatly along a *predatory-not predatory* binary, but instead exist along a spectrum of problematic and recommended publishing practices (Amsen, 2024; Nicholas et al., 2023). Indeed, some scholars have proposed that “even large, established subscription journal publishers such as Elsevier are predatory due to a perception of high subscription process and aggressive commercial behavior” (Linacre, 2022). These nuances and complexities may further complicate journalists’ decisions about what research to trust. Moreover, predatory journals’ tactics appear to be becoming more sophisticated over time (IAP, 2022), and will likely continue to do so as generative AI becomes more widely accessible. All of this points to the importance of evaluating “content” rather than “containers”—of scrutinizing the quality of the research rather than the reputation of the journal it is published in (Eve & Priego, 2017). However, our results suggest that journalists often start with the container before considering the content, at least among the experienced health, science, and environmental journalists we interviewed.

Journalists’ focus on journal reputation and prestige also has implications from a source diversity perspective, increasing the likelihood that research from newer, lesser-known journals—such as regional journals, OA journals, and those published in the Global South—will remain hidden from public view. While journalists’ preferential coverage of top (often, closed access) journals from the Global North is well documented (Conrad, 1999; Fleerackers et al., 2024; Nguyen & Tran, 2019), we found evidence that this bias may be becoming further entrenched amid growing concerns about predatory publishing and related research integrity issues such as hype, AI, fraud, and misinformation. Moreover, although there were exceptions, the journalists we interviewed appeared to be largely unaware, and sometimes unconcerned, about how their vetting strategies might perpetuate existing asymmetries in science media coverage (Nguyen & Tran, 2019). This has important implications for practice, as health and science journalists are actively working to diversify the sources they include in their coverage to present a more accurate, representative picture of the world (Newsome, 2021; Ordway et al., 2023). Journalists’ strategy of relying on “mainstream” journals exposes audiences to only a narrow swath of available research, potentially limiting their access to evidence that could help them make more informed choices. In light of these findings, communication scholars and educators may wish to consider ways to help journalists better recognize and understand predatory journals.

Our findings also have implications for scholarship, providing evidence into a topic that is timely and important but has been largely overlooked within the academic literature. Despite the important role that health, science, and environmental journalists play in brokering research knowledge to the public, they have not been identified as “key stakeholders” in global discussions about predatory publishing (IAP, 2022). Yet, those same discussions also note there is potential for predatory practices to erode public trust in academic research and damage public policy (*Ibid.*). Our study points to the importance of considering journalists in these discussions, as core players in whether and how the public is exposed to predatory publishing.

## Limitations

Importantly, our findings do not shed light on how journalists vet research *articles* but rather on how they evaluate the trustworthiness of the *journals* they are published in. Indeed, during their interviews, many journalists detailed rigorous strategies for assessing research article quality or credibility, including scrutinizing the methods or soliciting input from qualified independent experts. These vetting strategies were not analyzed in the present study but may help to mitigate some of the biases and limitations of the journal evaluation practices described above. Further research is needed to better understand these practices and their implications for the accuracy and diversity of the research that makes headlines.

Our results are also limited by our convenience sampling approach. Our use of professional science journalism groups and networks to recruit participants may explain why most journalists in our sample were highly experienced in reporting on health, science, and environmental topics. Moreover, all participants had some level of formal education, with many holding advanced degrees in STEM topics. These journalists likely offer an optimistic picture of journalists’ awareness of and approach to predatory journals, as less specialized journalists may be even less familiar with these journals and may not yet have developed strategies for avoiding them. In addition, our reliance on English-speaking journalists meant that most participants were based in North America. Future research is needed to understand how journalists working in different geographic, cultural, and linguistic contexts perceive and navigate predatory publishing practices. Doing so is essential for uncovering whether biases toward research produced and published in the Global North are also seen among journalists working in the Global South, as is often the case among scholars themselves (Krawczyk & Kulczycki, 2021).

## Author Contributions

Alice Fleerackers: Conceptualization, Formal Analysis, Methodology, Writing – Original Draft, Writing – Reviewing and Editing. Laura Moorhead: Conceptualization, Investigation, Methodology, Project Administration, Supervision, Validation, Writing – Original Draft, Writing – Reviewing and Editing. Juan Pablo Alperin: Validation, Writing – Reviewing and Editing.

## Acknowledgements

The authors wish to acknowledge Kyran Berlin for their research assistance. We also thank the members of the Scholarly Communications Lab who provided us with invaluable readings and context about the predatory publishing landscape, as well as the Journal Integrity Initiative research team for sharing insights that informed our research objectives and study design.

## Funding

This research was funded by an anonymous donor to Stanford University. AF is funded by a Social Sciences and Humanities Research Council of Canada (SSHRC) postdoctoral research fellowship (# 756–2023–0114).

### Data Availability

Due to the sensitive nature of the topic, data are only available upon request. The semi-structured interview protocol is available at: Moorhead, L. L., & Fleerackers, A. (2024). How journalists vet research and journals [Interview Protocol]. *OSF*. https://doi.org/10.17605/OSF.IO/WRTC4

